# Age and lead configuration matter: A comparative study of RF-induced heating of epicardial and endocardial electronic devices in adult and pediatric anthropomorphic phantoms in 1.5 T MR

**DOI:** 10.1101/2022.11.03.515092

**Authors:** Fuchang Jiang, Kaylee R. Henry, Bhumi Bhusal, Pia Sanpitak, Gregory Webster, Andrada Popescu, Giorgio Bonmassar, Christina Laternser, Daniel Kim, Laleh Golestanirad

## Abstract

**Background:** Children with congenital heart defects often have life-sustaining indications for a cardiac implantable electronic device (CIED). In children, these devices are typically sewn to the heart epicardium, but the FDA has never licensed an epicardial system as MR-Conditional due to limited data. Children’s hospitals default to either refusing MRI service to a vast majority of pediatric CIED patients or adopting a scan-all strategy based on results from adult studies. We argue that both approaches are flawed, and the risk-benefit decisions should be made on an individual basis.

**Purpose:** To provide evidence-based knowledge on RF-induced heating of CIEDs in children and adults with epicardial and endocardial leads of different lengths.

**Study Type:** Phantom

**Field Strength/Sequence:** 1.5 T.

**Assessment:** 120 clinically relevant epicardial and endocardial device configurations were implemented in adult and pediatric anthropomorphic phantoms. Temperature rise was recorded during RF exposure at 1.5 T.

**Statistical Tests:** Means comparisons were implemented using two-sample t-tests, reliability analysis using interclass correlation coefficient based on a single rating, absolute-agreement, 2-way mixed-effects model.

**Results:** There was significantly higher RF heating of epicardial leads compared to endocardial leads in the pediatric phantom (3.4 ± 3.0 vs. 0.6 ± 0.4 °C, p<0.001); however, there was no significant difference in the adult phantom (3.0 ± 3.2 vs. 2.0 ± 1.8, p=0.16). Endocardial leads in the pediatric phantom generated significantly less RF heating than in the adult phantom (0.6 ± 0.4 °C vs. 2.0 ± 1.8 °C, p<0.001).

**Data Conclusion:** Body size and lead length significantly affected RF heating. For models based on younger children with short epicardial leads (e.g., 25cm), RF heating up to 12 °C was observed, delivering a cumulative thermal dose previously associated with tissue necrosis. In contrast, RF heating in model based on children with endocardial leads was well below the heating expected from physiologic fever (3 °C).

## Introduction

Cardiac implantable electronic devices (CIEDs) treat cardiac conduction disorders, replacing sinus and atrioventricular nodal function and providing defibrillation for abnormal heart rhythms in adults and children. The common practice for implanting CIEDs in adults and older children (>10-25 kg) is to insert the leads through the subclavian vein and connect them to an implantable pulse generator (IPG) placed in the subpectoral pocket (1) (see Figure 1). In contrast, infants and young children often receive an epicardial CIED system, where the lead is sewn directly to the myocardium and the IPG is placed inferior to the abdominal rectus. While these conventions are most common in clinical practice, there are also cases where young children have received transvenous pacing systems and adult patients have received epicardial leads due to unusual venous anatomy (1).

**Figure 1:**
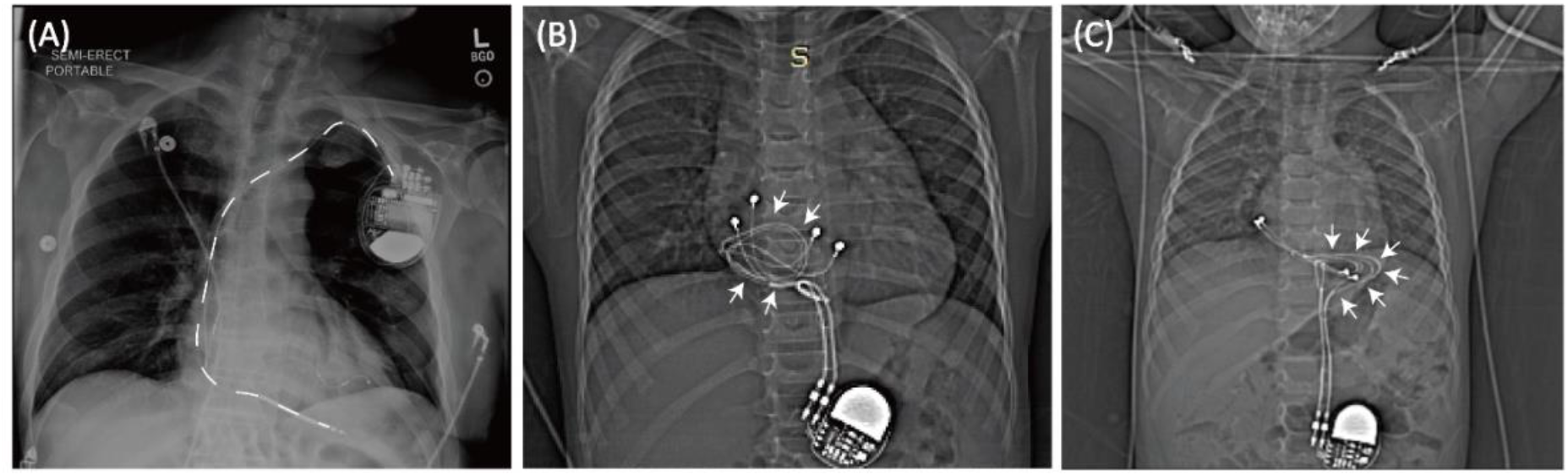
(A) Typical trajectory of an endocardial lead. Leads pass through subclavian vein creating a stereotypical pathway in nearly all patients. In contrast, epicardial devices can have highly varied lead trajectories (B-C).

Since 2011, MR-Conditional endocardial CIEDs have been available for adults, allowing patients to receive MRI exams under conditions that assure safety. In contrast, the FDA has never granted MR-Conditional status for an epicardial device and the limited data that is available indicates an elevated risk of RF heating of epicardial leads (2,3). To exacerbate the problem, a straightforward method to extract epicardial leads does not exist, and children who receive them may become ineligible to receive future MRI scans even if an MR-Conditional device is implanted later on.

The barriers to MRI in patients with epicardial leads are concerning considering that 75% of patients with CIEDs are indicated for MRI exams at least once during their lifetime (4) with many requiring repeated examinations (5,6). The demand is likely even higher in children, as developmental changes warrant more frequent assessments and alternative imaging modalities with ionizing radiations are more restricted.

The Heart and Rhythm Society’s 2017 expert consensus statement recommended that patients with CIEDs only undergo MRI when the product labeling is adhered to (3). The dire need for MRI, however, has led some groups to scan patients with epicardial systems off label (7,8). Reflecting this reality, the 2021 PACES guideline made a class IIb recommendation for pediatric CIED MRI patients with epicardial leads on individualized consideration of the risk/benefit ratio (9). However, no guidance is available on how to quantify those risks and the decision whether to scan a patient and the MRI protocol choice is based on the clinicians’ risks assessment which in turn is based on limited literature.

The goal of this study was to compare MRI-induced RF heating around the tips of epicardial and endocardial leads across varying lead lengths and trajectories inside adult and pediatric anthropomorphic phantoms during MRI at 1.5 T. Such a comparative study is currently missing in the literature and is necessary to allow evidence-based risks assessment.

Because body size and lead trajectory are two factors that substantially affect RF heating (10-16), we performed experiments in human-shaped phantoms of different sizes implanted with leads that followed patient-specific trajectories. To ensure our results were robust and reproducible, we repeated experiments for a subset of cases and performed a test-retest reliability analysis. Finally, to provide a conservative framework that is relevant to clinical care, we performed experiments to determine imaging conditions that constrained RF heating to < 3 °C (similar to a fever). Our work provides the knowledge on RF heating of epicardial leads in clinically relevant configurations as the basis for evidence-based assessment of risk-to-benefit ratio of performing MRI in children with CIEDs.

## Materials and Methods

### Phantom design and construction

Most studies that have assessed RF heating of conductive implants in the MRI environment have been performed in a box-shaped phantom inspired by the ASTM standards (17). Recently, however, a 10-fold discrepancy in the calculated specific absorption rate (SAR) in the ASTM box versus human body models was reported (18). This discrepancy was mainly attributed to the highly elliptical field polarization in the shallow-depth (∼9 cm) ASTM phantom, which substantially differs from the field distribution in the human body. For this reason, we fabricated custom-made human-shaped phantoms that mimicked the silhouette of an average-sized adult (length 65 cm, width 45 cm, depth 14 cm) and a twenty-nine-month-old child (length 53 cm, width 30 cm, depth 15 cm) (see Figure 2). Phantoms were filled with polyacrylamide (PAA) gel (22 L for the adult and 9 L for the pediatric phantom) with a conductivity of *σ* = 0.47 S/m and relative permittivity of *ε*_*r*_ = 88, representing dielectric properties of a tissue-mimicking medium. Attention was paid to ensure that implanted leads and the IPG were submerged within the gel at a depth of 2-3 cm below the surface, to mimic the placement in patients.

**Figure 2:**
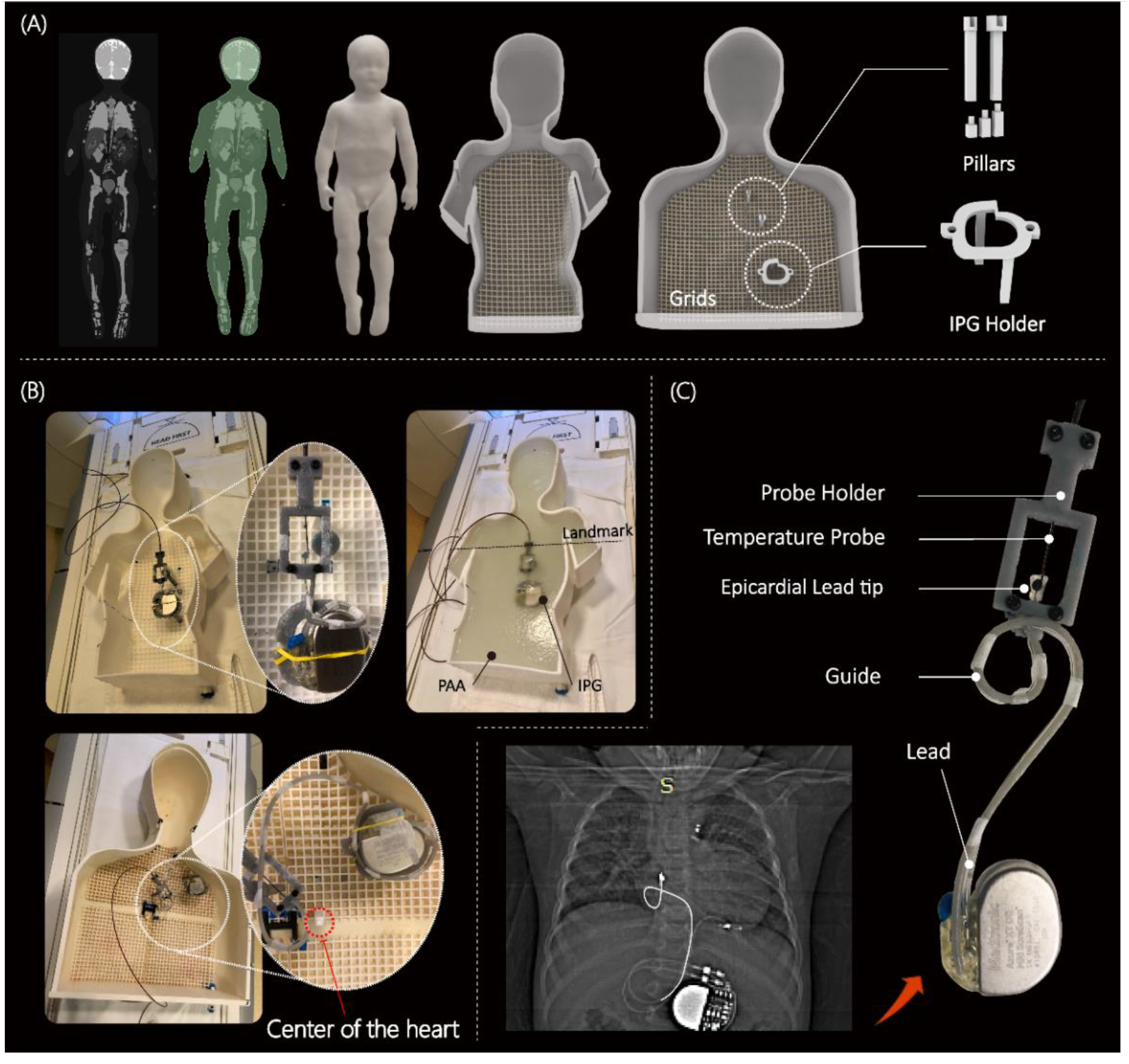
(A) MRI of a 29-month-old child was used to create 3D models of the child’s silhouette; Pediatric and adult phantom with girds, pillars, and different height of pillar extensions to adjust the positioning of the CIEDs. (B) Assembled phantoms with CIED and temperature measuring setup. Chest imaging landmark was used during MR scans. (C) An example of a trajectory derived from a patient x-ray image; Close view of contact between temperature probe and epicardial lead tip.

### Device configurations

RF-induced currents on elongated implants depend on the magnitude, polarization, and phase distribution of MRI incident electric field along the length of the implant (19-24). This means that the trajectory and position of an implanted lead has a decisive effect on the magnitude of RF heating around its tip (13,24-26). For this reason, a robust and reliable RF safety evaluation should include realistic patient-derived device configurations. We reviewed chest X-ray and computed tomography (CT) images of 200 adult and pediatric patients with epicardial and endocardial CIEDs to create representative device trajectories. Retrospective use of patient imaging data for modeling and safety assessment purposes was approved by the Institutional Review Boards of Northwestern University and Ann & Robert H. Lurie Children’s Hospital.

To enhance reproducibility, we designed and 3D printed trajectory guides. These helped route the leads along patient-specific trajectories and kept them securely in place during the experiments like our previous works (27,28) (see Figure 2C). Experiments with the epicardial systems were performed using the three of the most used lead lengths (i.e., 15 cm, 25 cm, and 35 cm) (Medtronic CapSure® EPI 4965). Leads were connected to a Medtronic Azure™ XT DR MRI SureScan pulse generator which was placed in the phantom’s abdomen. Experiments with the endocardial system were performed using 35 cm, 45 cm, and 58 cm leads (CapSureFix Novus MRI™ SureScan™ bipolar 5076). These were connected to the same pulse generator but placed in the left infraclavicular region.

Clinically relevant trajectories were modeled based on images of specific patients, X-ray images of patients available in the literature, or based on expert opinion of clinically important trajectories (authors GW, MM). A total of 60 experiments were performed in the child phantom, with 15-cm and 25-cm epicardial leads and 35-cm and 45-cm endocardial leads each routed along 15 patient-derived trajectories. Similarly, 60 experiments were performed in the adult phantom with 25-cm and 35-cm epicardial leads and 45-cm and 58-cm endocardial leads, each routed along 15 patient-derived trajectories. Figure 3 shows a few representative trajectories for each category. Computer-Aided Design (CAD) files for all 120 trajectories are available upon written request to the corresponding author.

**Figure 3:**
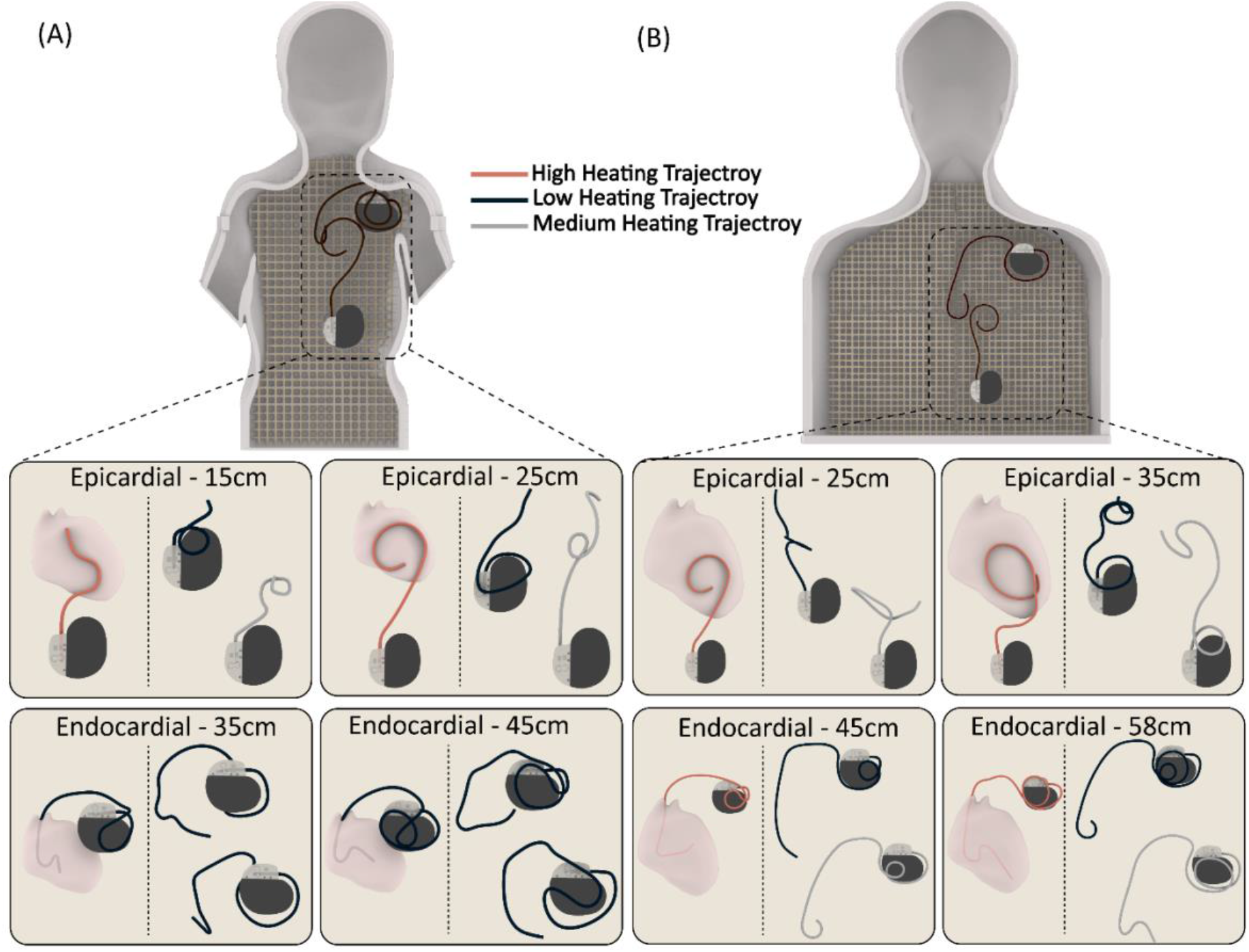
Three representative trajectories for each category (24 trajectories in total).

### Imaging protocol

Temperature measurements were performed using MR-compatible fiber optic probes (OSENSA, Vancouver BC, Canada, resolution 0.01 °C) secured at the tip of the lead. To ensure reliable thermal contact, we 3D printed a custom-designed holder that securely held the temperature probe in place so that it was in direct contact with the tip of the lead throughout the experiment.

RF exposure was performed in a 1.5 T Siemens Aera scanner (Siemens Healthineers, Erlangen, Germany) with the phantoms positioned such that the chest was at the isocenter, and a high-SAR T1-weighted turbo spin echo (T1-TSE) sequence (TE = 7.3 ms, TR = 897 ms, B_1_^+^ = 5 µT, acquisition time = 280 s). Although the acquisition time was shorter than what is commonly used in RF heating experiments (i.e., ∼5 minutes as opposed to 10 minutes) it was sufficiently long such that all temperature profiles reached the plateau, ensuring that our temperature rise comparative study rise was accurate.

### Test-retest analysis

To assess the reliability of our measurements, we repeated the experiments for 20% of the total number of cases (i.e., 24 re-test experiments) and calculated the Intraclass Correlation Coefficient (ICC). Repeated experiments were performed for trajectories that generated high, average, and low RF heating in each category, thus maximizing the heterogeneity of the re-test sample. ICC estimates and their 95% confidence intervals were calculated using RStudio based on a single rating, absolute-agreement, two-way mixed-effects model (29).

### Establishing safe exposure limits based on worst-case scenarios

Once comparative experiments were completed and validated, device configurations that generated the highest heating were identified for each category. Then, we performed secondary experiments to determine the maximum RMS B_1_^+^ that generated less than 3 °C RF heating after 10 minutes of scanning. To do this, we noted the device configuration that generated the worst-case RF heating in each category and implanted them into the phantom. We recorded the temperature at the lead’s tip during 10 minutes of scanning with varying RMS B_1_^+^ values. To have full control over the characteristics of the RF exposure, gradient coils were disabled and a train of 1 ms rectangular RF pulses was transmitted using the “rf_pulse” sequence from the Siemens Service Sequence directory. The flip angle was then adjusted to generate RMS B_1_^+^ values ranging from 2 µT to 5 µT. Pulse sequence parameters are summarized in Table 1 for each experiment. The upper limit (5 µT) corresponded to the scanner-reported SAR of 100%, which was the maximum RMS B_1_^+^ allowed before the scanner stopped the operation.

**Table 1:** Characteristics of the RF pulses.

|  |  |  |  |  |
| --- | --- | --- | --- | --- |
| Frequency<br>64MHz |  | Duration<br>603s |  | TR<br>24.03ms |
| Patient weight entered<br>68.0kg |  |  | Patient weight entered<br>13.6kg |  |
| Flip Angle (°) | B1+rms[μT] | Flip Angle (°) | B1+rms[μT] |  |
| 360 | 4.80 | 360 | 4.82 |  |
| 330 | 4.40 | 330 | 4.41 |  |
| 300 | 4.00 | 295 | 3.95 |  |
| 270 | 3.60 | 285 | 3.81 |  |
| 240 | 3.20 | 278 | 3.72 |  |
| 210 | 2.80 | 270 | 3.61 |  |
| 180 | 2.40 | 265 | 3.55 |  |
| 170 | 2.27 | 245 | 3.28 |  |
| 150 | 2.00 | 235 | 3.15 |  |
|  |  | 225 | 3.01 |  |
|  |  | 205 | 2.74 |  |
|  |  | 195 | 2.61 |  |
|  |  | 185 | 2.48 |  |
|  |  | 175 | 2.34 |  |

### Statistical Analysis

A means comparison was implemented using t-tests to determine differences in heating between epicardial and endocardial leads in the adult and child phantoms. All statistical tests were run using in RStudio (version 4.1.1) with a 95% confidence level.

## Results

### Reliability of measurements

The 95% confidence interval for the ICC was 0.95-0.99 indicating excellent reliability. RF heating measurement values for test-retest experiments along with corresponding device configurations are given in the Supplementary Information.

### RF heating of CIEDs in pediatric phantom

The RF heating mean ± standard deviation (SD) in the pediatric phantom was 0.6 ± 0.4 °C for endocardial leads and 3.4 ± 3.0 °C for epicardial leads (pooled over both lead lengths in each case). The maximum RF heating was recorded to be ∼12 °C for the 25-cm epicardial lead. For a 10-minute scan, this will be equivalent to a cumulative thermal dose of CEM43°C=1280 minutes, high enough to cause necrosis in pig muscles (30).

A one-tail t-test showed a significantly higher epicardial lead’s RF heating compared to endocardial leads in the pediatric phantom (p<0.001). Within epicardial leads, the 25-cm lead generated higher RF heating than the 15-cm lead (4.4 ± 3.7 °C vs. 2.4 ± 1.5 °C, p<0.05). Within endocardial leads, the 35-cm lead generated higher RF heating than the 45-cm lead (0.9 ± 0.4 °C vs. 0.4 ± 0.3 °C, p<0.001).

### RF heating of CIEDs in adult phantom

The RF heating’s mean ± standard deviation in the adult phantom was 2.0 ± 1.8 °C for endocardial leads and 3.0 ± 3.2 °C for epicardial leads. There was no significant difference between RF heating of endocardial and epicardial leads in the adult phantom (p=0.16). The maximum RF heating was ∼12 °C occurring for the 25-cm epicardial lead. Within epicardial leads, the 25-cm lead generated higher heating than the 35-cm lead (4.8 ± 3.5 °C vs. 1.2 ± 1.5 °C, p<0.001). Within endocardial leads, 45-cm lead generated higher heating than the 58-cm lead (3.2 ± 1.9 °C vs. 0.8 ± 0.7 °C, p<0.001).

### Adult vs. pediatric phantom

There was no significant difference between RF heating of epicardial leads in the pediatric and adult phantom (p=0.65). However, endocardial leads in the pediatric phantom generated significantly less RF heating compared to endocardial leads in the adult phantom (0.6 ± 0.4 °C vs. 2.0 ± 1.8 °C, p<0.001). This could be attributed to two factors. First, the incident MRI electric field tends to be lower in a child’s body compared to an adult body because of a child’s smaller size and a more central position within the RF coil. This can be observed in Figure 5. Figure 5 shows the simulated MRI incident E field (i.e., electric field in the body in the implant’s absence) in a 29-months-old body model compared to an adult body model, subject to RF exposure in a 64 MHz body coil. The input power of the coil was adjusted to generate a mean B_1_^+^ = 5 μT at the isocenter for both. CIED devices would be exposed to a higher incident electric field in the adult body compared to the child body (Details of simulation setup and results are given in the Supplementary Information).

The second factor is attributable to differences in lead trajectories related to children’s anatomy. Because the IPG is much closer to the heart in pediatric patients compared to adults, there will be more excess lead length, which is usually looped around the IPG. The addition of these loops could have contributed to the lower RF heating similar to what is observed in neuromodulation devices (31,32). This can be specifically appreciated comparing RF heating of the 45-cm endocardial leads in adult (3.2 ± 1.9 °C) vs. child (0.4 ± 0.3 °C) phantoms, where the difference in RF heating can be solely attributed to differences in position and trajectory of the lead (Figure 4).

**Figure 4:**
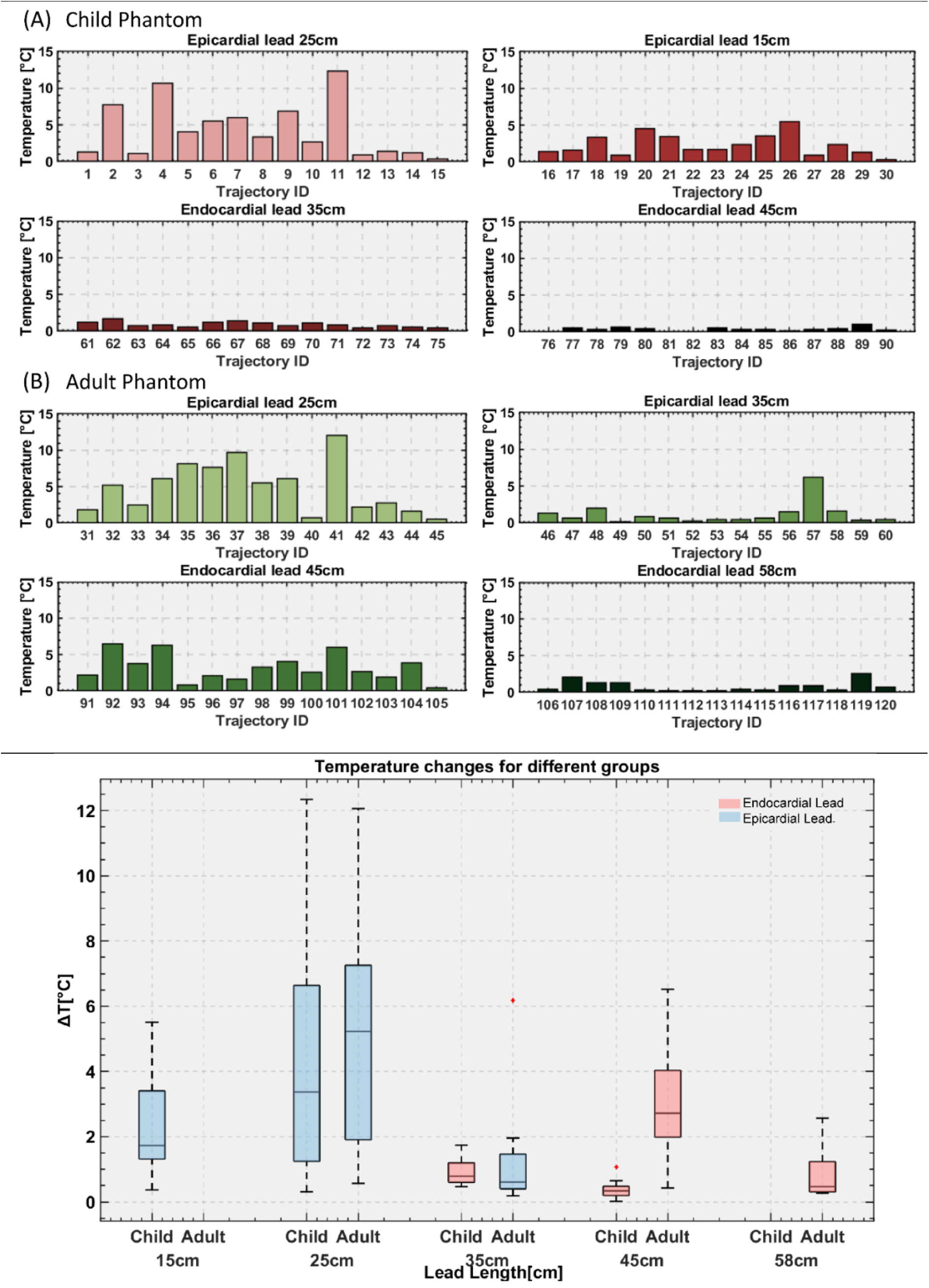
Temperature rise at tips of the lead with 120 different trajectories in pediatric and adult phantoms.

### Effect of lead’s length

The maximum RF heating in both phantoms was observed for the 25-cm epicardial lead. The half-wavelength of radiofrequency field in the PAA (*εr*= 88) was ∼ 24.5 cm, which explains the high heating as the length of the lead approached the resonance length. This phenomenon, commonly referred to as the resonance effect, has also been observed in other types of wire implants (33,34).

### Estimation of safe RMS B_1_^+^ levels based on worst-case observations

Temperature rise distribution histograms are given in the Supplementary Information. For each case, the device configuration that generated the highest RF heating was used in subsequent experiments to estimate RMS B_1_^+^ values that constrained RF heating to < 3 °C. Our data indicates that this approach was conservative, as > 99% of cases in the general population would generate RF heating below this limit.

Figure 6 gives the RMS B_1_^+^ values and their corresponding RF heating for device configurations that generated highest heating in each category. For the pediatric phantom, both 35-cm and 45-cm endocardial leads generated RF heating well below 3 °C after 10 minutes of scanning at the maximum allowable RMS B_1_^+^ (i.e., 5µT) and thus, are not shown in the graph. The maximum RMS B_1_^+^ that generated RF heating < 3 °C was 2.5µT for the 25-cm epicardial lead and 3.6µT for the 15-cm epicardial lead.

**Figure 5:**
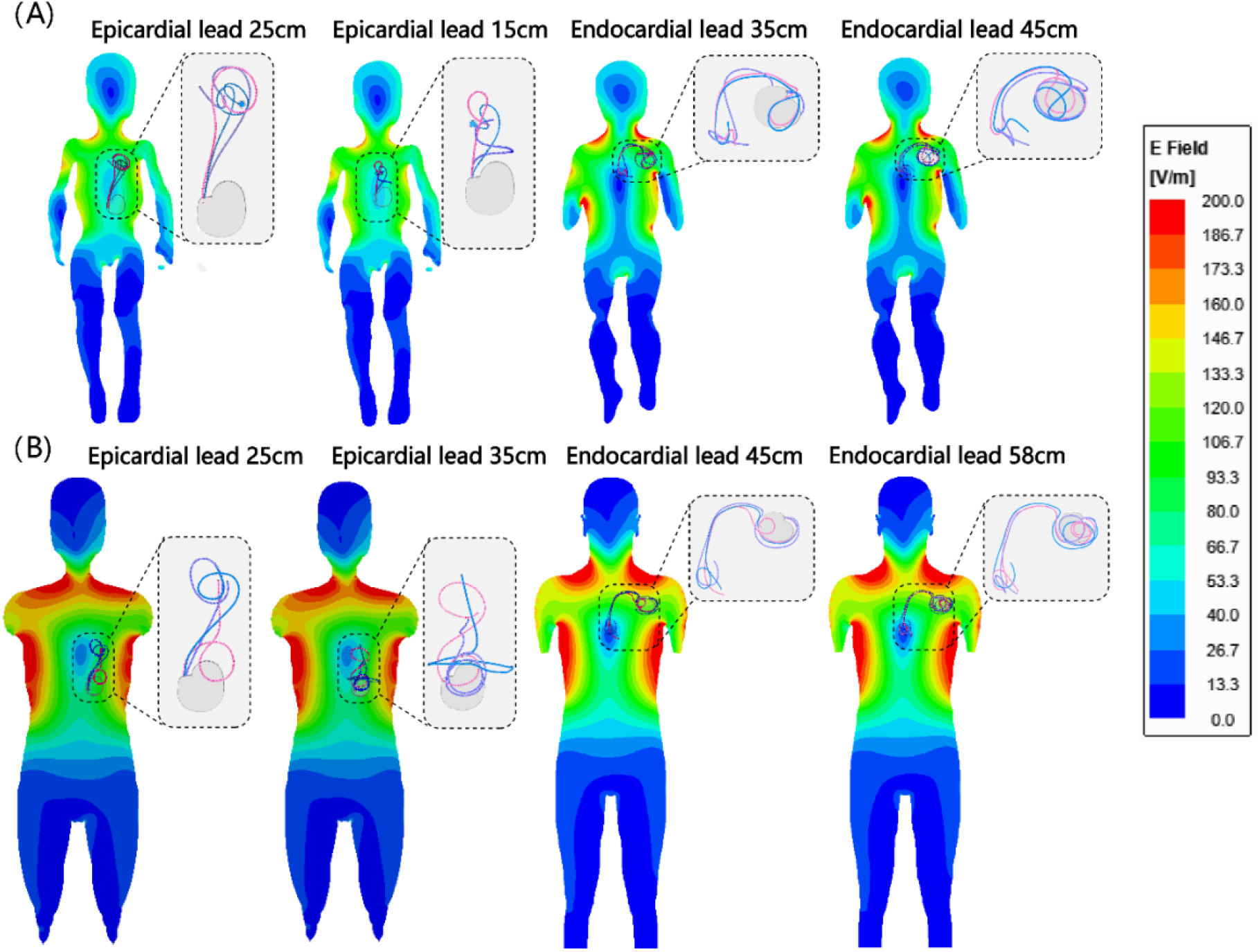
Simulated MRI incident E field for both homogeneous child body model (A) and adult body model (B) with three superimposed endocardial and epicardial device configurations for each case (24 configurations in total).

**Figure 6:**
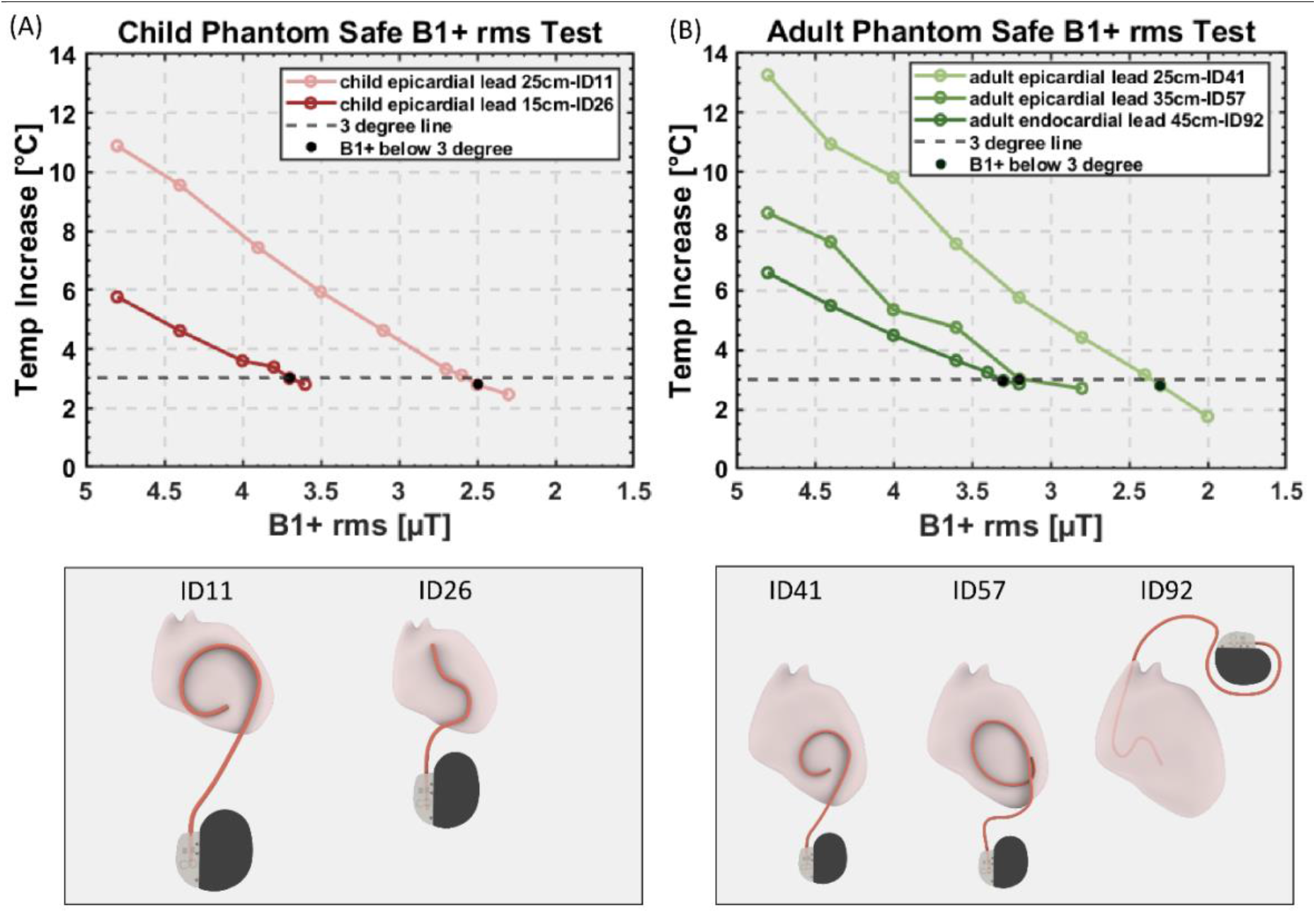
B_1_^+^ RMS values and their corresponding RF heating for device configurations that generated highest heating in each category. Black dots: B_1_^+^ RMS value that generates the temperature rise below 3°C.

For the adult phantom, the 58-cm endocardial lead generated RF heating well below 3 °C so it is not shown in the graph. The maximum RMS B_1_^+^ that generated RF heating < 3 °C was 2.3µT and 3.2µT for the 25-cm and 35cm epicardial lead, respectively and 3.3µT for the 45-cm endocardial lead.

## Discussion

### Risks of MRI in patients with CIED

An MRI machine produces three electromagnetic fields which can interfere with a CIED: the static magnetic field (B_0_), the RF field (B_1_ and its associated E field), and the pulsed magnetic gradient field (i.e., G_x_, G_y_, G_z_). The interaction with B_0_ is negligible in the modern era. Gradient fields can induce electric currents in the device’s circuitry potentially interfering with device function, however, new generation of CIEDs have enhanced programming and fail-safe MRI switches that minimize the pulsed magnetic gradient field interference. RF heating, on the other hand, remains a major issue. This occurs due to the “antenna” effect, where the transmit coil’s electric field couples with CIED leads and amplifies the energy deposition in the tissue at the tip of the lead, causing potential thermal injuries.

Two interplaying factors determine RF heating severity for implanted CIEDs: patient’s body size and the lead’s length/ trajectory. Patient body size is important because the MRI electric field’s distribution inside a sample is highly sensitive to the sample’s size and material. The different body size and material proportion in children (e.g., bone, fat, and muscle ratio) interacts with the MRI field differently than adult body composition. This leads to a substantially different electric field distribution at the CIED’s location. For this reason, MRI safety assessment for children must be performed in age-specific pediatric body models.

The second determinant of RF heating is the lead trajectory which is correlated with patient size; smaller children receive shorter leads. It is important to note that certain lead lengths that are common for adults and children typically follow a different trajectory in children (e.g., more loops around the IPG to account for the excess length). Because the position, orientation, number of loops, and lead length within the human body determine the degree to which MRI electric field couples with the lead, RF safety in children must be assessed on an age-specific basis. Our data shows that certain lead configurations in children generate RF heating that is an order of magnitude lower than what FDA deems to be safe in adults (e.g, 35-cm and 45-cm endocardial leads). However, some epicardial lead configurations generate excessive heating high enough to cause myocardial lesions within 10 minutes (e.g., 25-cm epicardial lead), highlighting the need for risk assessment specific to size and lead configuration.

### MRI in Children with CIEDs

There are Class I indications for CIED therapy in children with congenital heart defects (CHD), inherited arrhythmia syndromes, and congenital disorders of cardiac conduction (9). Unlike adult CIED patients, there exists a large registry data totaling several thousand patients that demonstrate MR safety at 1.5T, MR safety. These data are severely lacking for pediatric CIED patients. Since CIED lead types and IPG configurations are substantially different in children compared to adults, it is important to provide more research on this topic; a simple transference of adult results for children is inappropriate. The 2017 HRS consensus statement highlights this knowledge gap, “given the paucity of data related to the safety of MRI [with epicardial leads], recommendations cannot be made [and] many questions remain unanswered.”(3) Despite this lack of evidence, the 2021 PACE guideline made class IIb recommendation for pediatric CIED MRI patients with epicardial or abandoned leads on an individualized consideration of the risk/benefit ratio (9). However, the 2021 PACE guideline did not provide specifics as how to quantify those risks. Since individualized risk-benefit decisions are hard to make without data, children’s hospitals default to either refusing MRI service to most pediatric CIED patients. Alternatively, children’s hospitals adopt a scan-all strategy based on results from adult studies. We argue that both approaches are flawed. A blanket ban on MRI means that children are referred for alternative imaging modalities with ionizing radiation, which not only increases the risk for cancer (35), but also results in suboptimal diagnosis, prognosis, and surgical planning. On the other hand, a scan-all strategy could expose some children to dangerously high RF heating as shown in this study. Our data thus highlights the need for age-specific assessment of RF heating in children with CIEDs.

### Rise of off label scanning is a reality and caution is warranted

So far, off label MRI of CIED patients have been performed in several thousand adults and reported no significant side effects. The word “significant”, however, should be interpreted with caution. A recent study on adults with abandoned or functioning epicardial leads (36) found a transient elevation of pacing threshold in one patient, and an irreversible atrial pacing lead impedance elevation (>10,000 ohms) in another. While these may have been coincidental to the timing of the MRI, the retrospective nature of the study and the timing of the impedance rise make RF heating in the MRI scanner impossible to exclude. It is possible that RF heating in the MRI scanner led to a formation of low-conductivity scar tissue around the tip of the lead. Another study in pediatric and adult patients with epicardial or abandoned leads (7) reported a heat sensation in the left lateral chest at the location of an abandoned endocardial lead in one patient (36 years old) — which required the scan to be stopped — and tingling sensation at the location of epicardial lead in another. Even if these risks may be manageable in adults, there are additional factors that complicate accurate risk assessment in children. An infant or a 5-year-old scanned under anesthesia cannot complain of a heating sensation to stop the MRI. The results could be an irreversible thermal injury of the heart tissue. Because myocardial scarring during early childhood can expand as the heart grows over time (37), such damages could cause long-term complications over the individual’s life span. While we did not find a systematic difference between the pediatric and adult phantoms, marked heating elevations were present in clinically-relevant scenarios. For example, our results show that MR-induced heating of epicardial leads reaches levels that are 267% higher in young children who mostly receive short epicardial leads (e.g., 25-cm) compared to adults who receive longer leads (e.g., 35-cm). This study is the first step to identifying characteristics of children who represent edge cases at elevated risk for tissue heating, making age-specific assessment of RF safety even more crucial.

### Limitations and future work

This work has several limitations. First, only a few epicardial and endocardial lead models were investigated. We are aware that other lead models exist in the market. Even within device models in our study, we limited our work to lead lengths that were most commonly used at our institution. For example, Medtronic 4965 epicardial leads also come in 50-cm length, and Medtronic 5076 endocardial leads also come in 52-cm, 65-cm and 85-cm lengths. Bipolar epicardial leads (e.g., Medtronic 4968 CapSure Epi) were not investigated either. This was a compromise that allowed us to study each lead more thoroughly (i.e., more patient-derived trajectories) which we deemed to be crucial to infer conclusions on safety.

Second, this work only studied intact CIED devices; abandoned leads were not included. Abandoned leads are observed in a sizeable fraction of CIED patients for whom leads fail for technical reasons, infection, or blockage of the vein by clot. In such situations the patient is referred for lead extraction and implantation of new leads (38) or lead abandonment and implantation of additional leads (39). Extraction is not always successful in transvenous leads and the risks of sternotomy and surgical lead extraction in patients with epicardial leads typically outweigh the benefits. This suggests that fractured epicardial leads are nearly always abandoned in place. Since MR-Conditional labeling of a CIED only applies to the intact device with leads implanted in their original configuration, patients with abandoned leads are contraindicated for MRI. A recent study showed that capped abandoned leads induce tissue heating that is on average 3.5 times higher than what is generated by a complete CIED system (40). Thus, results of this study should not be generalized to cases of abandoned leads.

