## Supplementary Information for "Age and lead configuration matter: A comparative study of RF-induced heating of epicardial and endocardial electronic devices in adult and pediatric anthropomorphic phantoms in 1.5 T MR"

Supplementary Information Figure S1

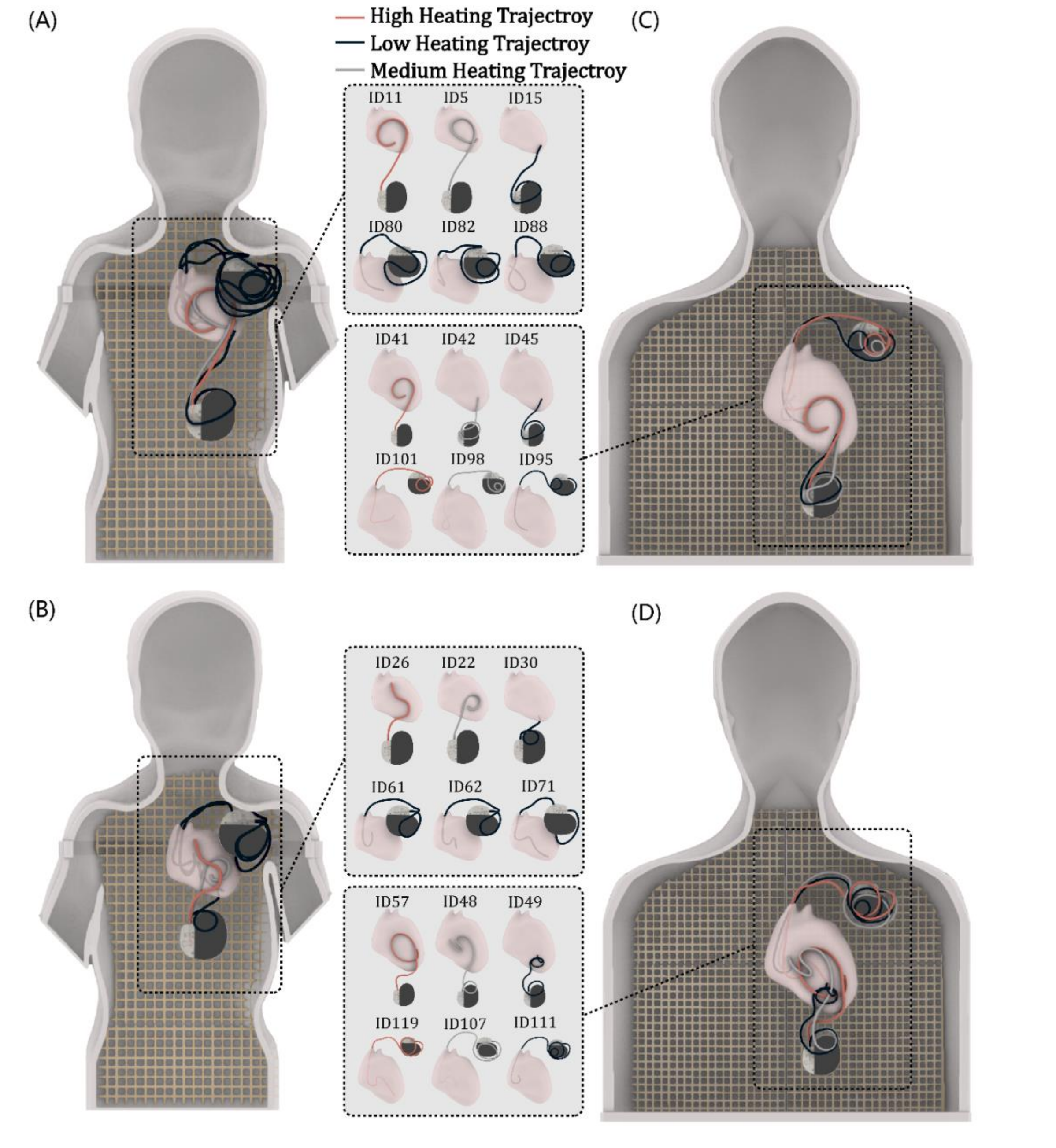


Figure 1: superimposed test-retest selected trajectories. (A) pediatric phantom with 25cm epicardial and 45-cm endocardial trajectories (6 in total), (B) pediatric phantom with 15-cm epicardial and 35-cm endocardial trajectories, (C) adult phantom with 25-cm epicardial and 45-cm endocardial trajectories and (D) adult phantom with 35-cm epicardial and 58-cm endocardial trajectories.

Supplementary Information Table S1

| Child Phantom | Reproduced temp[°C] | original temp[°C] | Adult Phantom | Reproduced temp[°C] | original temp[°C] |
|---|---|---|---|---|---|
| ID11 | 10.51 | 12.34 | ID41 | 13.06 | 12.06 |
| ID5 | 4.52 | 4.04 | ID42 | 1.41 | 2.18 |
| ID15 | 0.52 | 0.31 | ID45 | 0.57 | 0.28 |
| ID26 | 5.35 | 5.51 | ID57 | 5.86 | 6.18 |
| ID22 | 1.23 | 1.73 | ID48 | 1.2 | 1.98 |
| ID30 | 0.86 | 0.37 | ID49 | 0.27 | 0.19 |
| ID62 | 2.47 | 1.74 | ID101 | 5.97 | 5.90 |
| ID61 | 1.38 | 1.25 | ID98 | 4.58 | 3.22 |
| ID71 | 0.51 | 0.84 | ID95 | 2.42 | 0.8 |
| ID80 | 0.35 | 0.44 | ID119 | 2.43 | 2.57 |
| ID88 | 0.21 | 0.39 | ID107 | 1.53 | 2.05 |
| ID82 | 0.04 | 0.09 | ID111 | 0.37 | 0.27 |

Table 1: test-retest temperature increases for each trajectory.

Supplementary Information Figure S2

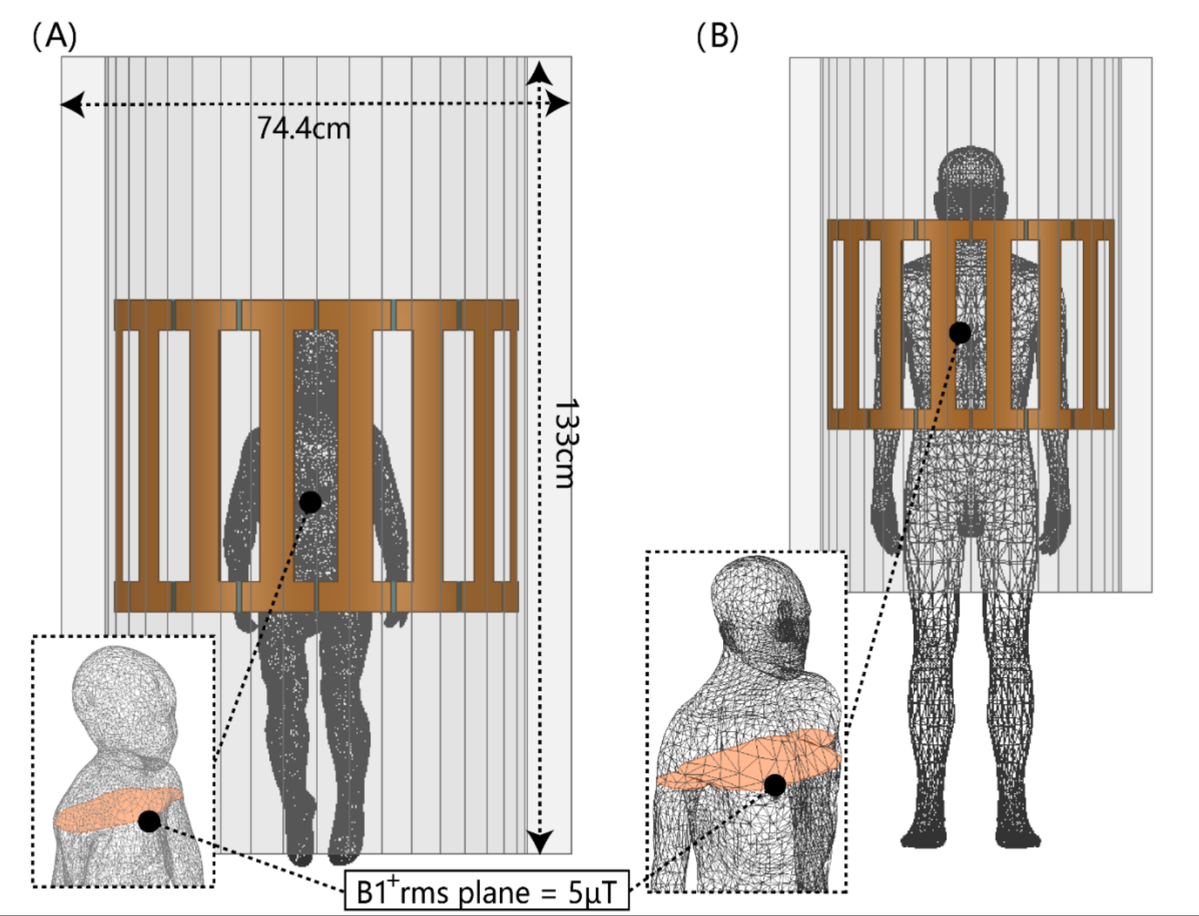


Figure 2: (A) The homogenous child body model (σ = 0.47 S/m; $\varepsilon_r$ = 80) placed inside a model of Siemens 1.5 T Aera body coil with the heart at the iso-center. (A) The homogenous adult body model placed inside a model of Siemens 1.5 T Aera body coil with the heart at the iso-center.

Supplementary Information Table S2

| Parts | Num Tets | Min edge length(mm) | Max edge length(mm) | RMS edge length(mm) | Min tet vol($mm^3$) | Max tet vol($mm^3$) | Mean tet vol($mm^3$) | Std Devn (vol)($mm^3$) |
|---|---|---|---|---|---|---|---|---|
| Child Body | 105328 | 0.46 | 26.22 | 14.88 | 9.72E-05 | 759.15 | 126.50 | 89.70 |
| Adult Body | 327401 | 2.36 | 22.89 | 16.04 | 1.83E-02 | 704.40 | 204.46 | 83.82 |

Table 2: Mesh statistics for the simulations for both child and adult.

Supplementary Information Figure S3

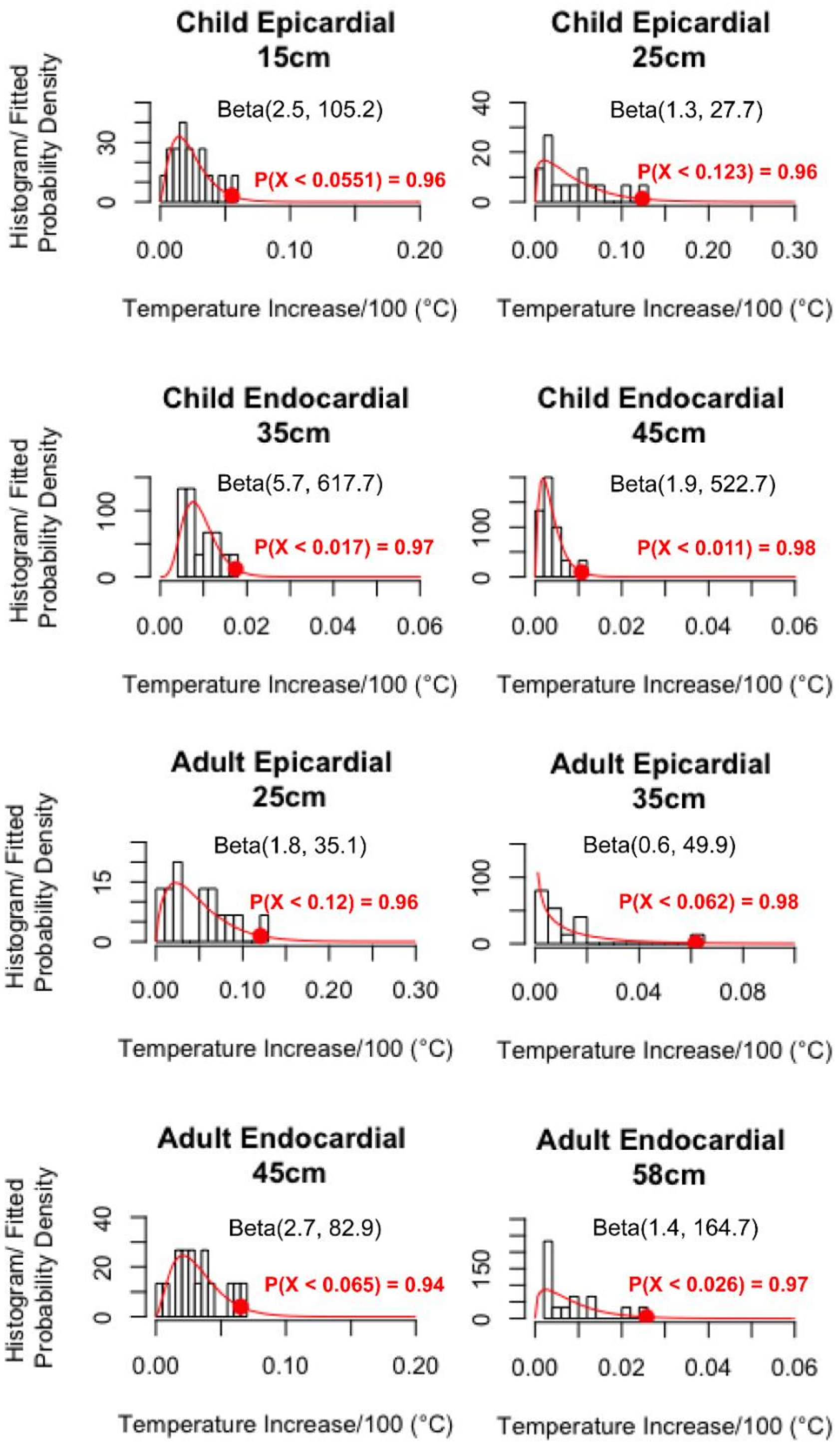


Figure 3: Scaled temperature increase histograms for both phantoms in each category. A beta probability density function was fitted to the temperature increase distribution for each case. The probability of a temperature increase occurring below the highest heating determined during the experimental setup is shown for each scenario.
